# Morphogenesis of a stratified cell mound at a vortex defect

**DOI:** 10.64898/2026.08.17.745181

**Authors:** Hsiang-Ying Chen, Carles Blanch-Mercader, Caroline Giuglaris, Jacques Prost, Pascal Silberzan

**Author notes:** Equal contribution (Hsiang-Ying Chen, Carles Blanch-Mercader).

## Abstract

Although topological defects in cell monolayers have been recognized as mechanical organizing centers in morphogenetic processes, the mechanism by which cells coordinate their motion at such defects and self-organize into higher-order structures remains elusive. Here, we report the formation of three-dimensional (3D) multicellular mounds in unconfined myoblast monolayers, at well-controlled vortex topological defects. Prior to the onset of bilayering, the vortex structure induces millimeter-scale cell flows converging toward the defect center. As a result, 3D cell mounds form at the defect core, layer-by-layer. These mounds grow by interlayer permeation sustained by the converging cell flows. At late stages, the bell shape of the structured mounds can be modeled with a dynamics driven by these converging flows. Our results therefore highlight the crucial role of integer topological defects in driving large-scale cell flows yielding the formation of highly ordered 3D tissues from a monolayer. We propose that similar mechanisms may be at play in certain morphogenetic and tumorigenic events.

---

Morphogenesis of tissues results from the interplay of physical forces and biochemical signals at all scales [1–4]. For instance, collective migration and pattern formation that shape the embryo [4–7] are driven by cell shape changes, intercalations, or divisions that are themselves mediated by signals including mechanical forces. Interestingly, the different tissues of the developing embryo are formed from cellular monolayers encapsulating a cell-free cyst [8]. Yet the formation of three-dimensional structures from a bidimensional cell monolayer remains an open question that also relates to the formation of 3D tumors from epithelia [9–11]. By “3D tissues”, we mean dense bulk assemblies of cells such as muscles, in contrast with hollow structures resulting from the wrapping or folding of a cell monolayer.

In the past decade, monolayers of spindle-shaped cells have been successfully described as active nematic liquid crystals [12–15], providing novel insights into developmental and pathological processes. Importantly, this description outlines the intrinsic presence of topological defects where stress patterns trigger individual [16] or collective [17,18] cell extrusion. For instance, in vitro, topological defects are the loci of formation of myoblast “crisscross bilayers” [18,19], 3D mounds of neural progenitor cells [17], and accumulation of fibroblasts [20–22]. In vivo, defects also appear critical in embryogenesis [23,24], morphogenesis [25–28], or tumorigenesis [29,30]. For instance, topological defects act as biochemical and mechanical organizing centers in the regeneration of Hydra [25]. Moreover, cell vortexes (also called “swirls”) are critical for proper cell alignment and differentiation in osteogenesis and play therefore a central role in the formation of organized and functional bone tissue [28]. By contrast, cell swirls are a criterion for diagnosing pathological situations, such as papillary thyroid carcinoma [31].

In vitro, topological +1 charges have been previously generated by tight spatial confinement in disks [27,32], giving rise to cell mounds at their center. Therefore, experiments where cell orientation is controlled as a +1 defect in a border-free monolayer are needed to unambiguously decipher the impact of these integer topological defects on the formation of 3D organized tissues, independently of confinement.

Here, we study the dynamics of myoblasts selected against differentiation plated on extended substrates textured with concentric circles amounting to a +1 vortex defect. When plated on featureless substrates, these cells organize in active nematic monolayers incorporating half-integer topological defects with 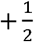 defects self-propelling “tail first” as previously observed [17,32]. When plated on vortex patterns, the cells align with the direction of the micro-rails and develop a millimeter-range convergent flow that results in collective cell extrusion and multilayer formation at the core. Eventually, cells form a 3D bell-shaped cell mound of complex architecture.

To understand these observations, we develop a hydrodynamic theory showing that a combination of active stresses, active nematic traction forces and orientation gradients generates long-range in-plane radial converging flows and short-range out-of-plane permeation flows, leading to layer-by-layer mound formation and, eventually, to the bell shape of the mound.

## 3D mound formation in a cell monolayer at a vortex defect

C2C12 mouse myoblasts were plated on polydimethylsiloxane (PDMS) extended substrates (13mm*13mm) whose surfaces were microstructured with concentric circular micro-rails, describing a +1 vortex defect (Methods, Extended data Fig. 1a-b) [20,22,33]. At confluence, cell monolayers reached a thickness of 6 ± 1 µm and exhibited an orientation field mirroring the vortex structure imposed by the underlying substrate (Extended data Fig. 1c-e, Extended data Fig. 2c). Starting approximately 12h after confluence, monolayers gradually developed a 3D multicellular mound at the defect core (Figure 1a, Extended data Fig. 2c, Supplementary video 1), reaching a height of 85 µm ± 16 µm (standard deviation, SD, n=20) 72 hours post-confluence (from now on, we take t = 0 h at confluence). Mound formation was exclusively observed at the defect core. These experiments being conducted on centimeter-scale patterned substrates, the monolayer remains unconfined, and we can conclude that integer defects are organizing centers for 3D mounds. Importantly, cells remained parallel to the substrate with negligible out-of-plane deviation. Indeed, the mean elevation angle *ϕ* was systematically smaller than 0.5° (Extended data Fig. 3). Initially (t < 30 h), the vortex orientation propagated across all superimposed cell layers that formed with time in a process mediated by cell-secreted extracellular matrix (Extended data Fig. 4d). Note that the standard deviation of *ϕ* is ~ 10 deg. So, cells explore adjacent layers but, on average, these out of plane contributions cancel out.

**Figure 1:**
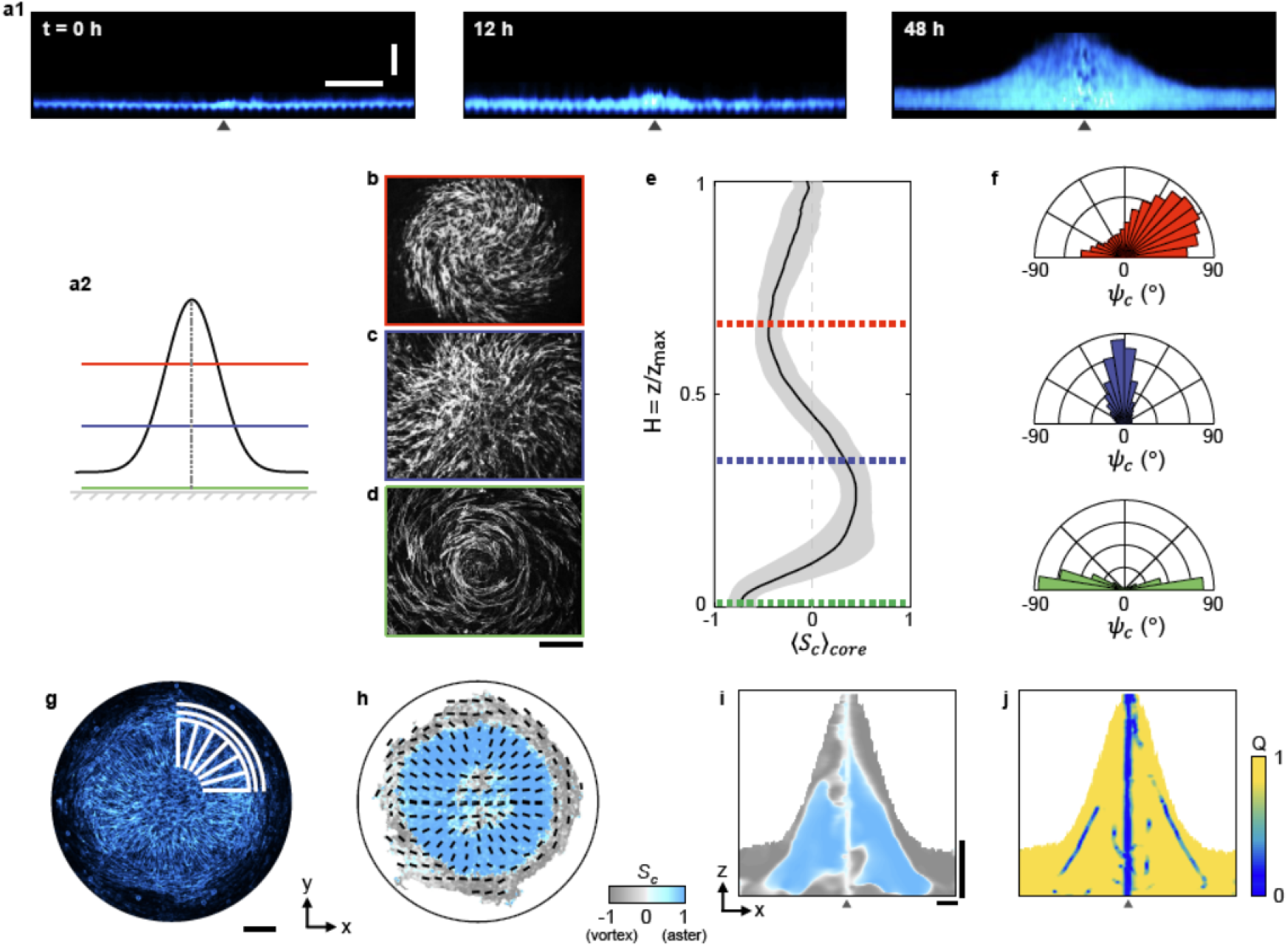
Complex nematic order in mature cell mounds. **a1**, Formation of a representative mound. Angle-averaged transverse sections ([xz] plane) along a diameter obtained from confocal imaging of a developing cell mound at t = 0 h, 12 h, and 48 h. C2C12 β-actin–mCherry myoblasts cells. The black triangle indicates the position of the core of the defect. See also Supplementary video 1. **a2**, Schematic of a late-stage cell mound. The coloured horizontal lines picture the positions of the representative confocal slices in panels b-d. **b-d**: Representative confocal images (topview, [xy] plane) of a mature cell mound fixed at t = 55 h for three values of the rescaled height = 0 (green), 1/3 (blue) and 2/3 (red) being the height coordinate, and the mound height. Note the organization of the cells as vortex, aster, and spiral according to the relative altitude in the mound. 5 % of the cells were C2C12 β-actin–mCherry cells. **e**: Order parameter being the angle between the radial direction and the director, averaged over a region 400 x 400 μm^2^ around the defect center. is plotted as a function of the rescaled mound height =z / z_max_. Dashed lines indicate the position of the confocal slices with the same color code as panels **b-d**. See also Extended data Fig. 1g, Extended data Fig. 5, and Supplementary video 2. Error bars are SDs. **f**: Probability density functions of *ψ*_*c*_ at various heights (same color code as panels **b-d**). Close to axis, the cells orient as vortex in the bottom layer (green), as aster in the middle layers (blue) and as spiral (angle = 52 °) in the top layers (red). **g**,**h**: Representative single confocal slice (*H* = 2/3) of a mature cell mound (**g**), and corresponding orientation field (**h**), showing the aster geometry within the mound (blue), and the vortex orientation at its edge (grey). Fixed actin-labelled cells. The colors encode the local value of *S*_*c*_. White lines in panel **g** are guides for the eye of the director field. Note the wider FOV of panel **g** compared to panels b-d. **i-j**: Representative [xz] cross-section of the order parameters *S*_*c*_ and *Q* in a mature cellular mound (t=65 h) (see Methods). Note the high degree of order in the mound (*Q*~1) except at the disclination line connecting the 2D defect core (*x* = *y* = *z* = 0) and the apex of the structure (no nematic order, *Q*~0). Unless otherwise specified, horizontal scale bars are 100 μm and vertical scale bars are 25 μm. Statistical information is provided in Extended data Table 1.

In a later regime (t > 30 h), the first layer kept the vortex orientation but the orientation field of the subsequent stacked layers changed to aster in most of the mound and to spiral in its top part (Figure 1b-f, Extended data Fig. 5, Supplementary video 2). In the following, multilayers are called “young mounds” when 12 h < t < 30 h, “developing mounds” for 30 h < t < 45 h, and “mature mounds” for t > 45 h.

We quantified the nematic order in mature cell mounds with the in-plane nematic order parameter *S*_*c*_ *=* cos(2*ψ*_*c*_) as a function of height (see Extended Data Figure 1 and Methods). *S*_*c*_ *=* − 1 (resp. +1) corresponds to a vortex (resp. aster) orientation. ⟨*S*_*c*_⟩_*core*_ is an average of *S*_*c*_ over a 400µm*400µm FOV centered at the core axis. Note that this FOV is smaller than the lateral extension of a mature mound. Variations of ⟨*S*_*c*_⟩_*core*_ along the *z*-axis showed three regions: a vortex orientation in the first layer (in direct contact with the microstructured substrate), an aster orientation in the middle layers and a spiral orientation in the upper layers of the mound (Extended data Fig. 5 and Extended data Fig. 6). In addition, when altitude was rescaled by the mound height, the profiles of ⟨*S*_*c*_⟩_*core*_ collapsed into a single master curve (Methods, Figure 1e and Extended data Fig. 6).

Of note, the +1 topological charge was preserved in all optical confocal slices spanning the height of mature mounds, thereby defining a line-defect where order is lost, perpendicular to the substrate plane and connecting the initial defect core to the apex of the mound (Figure 1j). This line corresponds to small values of the local nematic order parameter *Q* that quantifies the degree of nematic order [32] (see Methods) (Figure 1j).

The probability density functions of *ψ*_*c*_ at different heights quantified the above observations (Figure 1f) and showed that the angle of the spirals at the top of the mound peaked at 52° ± 17°. Interestingly, analyzing the orientation field further away from the mound axis showed that the aster central domain was enclosed by a vortex domain with a sharp transition zone (30 – 40 µm, comparable to a cell size) (Figure 1g-i).

To confirm that the apex of the mounds hosts a +1 topological defect and not a change of direction of the cells or a loss of their spindle shape, we measured their anisotropy in that region with a Shape Index parameter SI (Methods). We measured SI = 2.1 ± 0.7, indicating that cells remain well-elongated. For comparison, SI = 3.0 ± 0.8 in the vortex regions, reflecting the larger deformation of the cells.

## Long range convergent cell flows yield collective extrusion

We first investigated the initial global cell flows in the mound by analyzing phase contrast movies (Figure 2, Supplementary video 3). When reaching confluence (t = 0 h), myoblasts migrated actively along the vortex-structured micro-rails by contact guidance [34–36]. (Figure 2a-b, left panels, Supplementary video 3). After confluence, the azimuthal component of the velocity, ⟨*v*_*θ*_⟩ fluctuated around zero at all times, indicating the absence of a net rotational flow (Figure 2d). In parallel, a radial flow directed toward the defect core gradually developed with time (Figure 2a-d, Supplementary video 4). Myoblasts then extruded collectively at the defect core starting at t_e_ = 12 ± 2 h and organized as a bilayer in a first step of the formation of a mound (Figure 1a, Extended data Fig. 2c). Of note, in this convergent flow, cells kept their initial orientation and moved perpendicularly to their long axis (Supplementary video 5).

**Figure 2:**
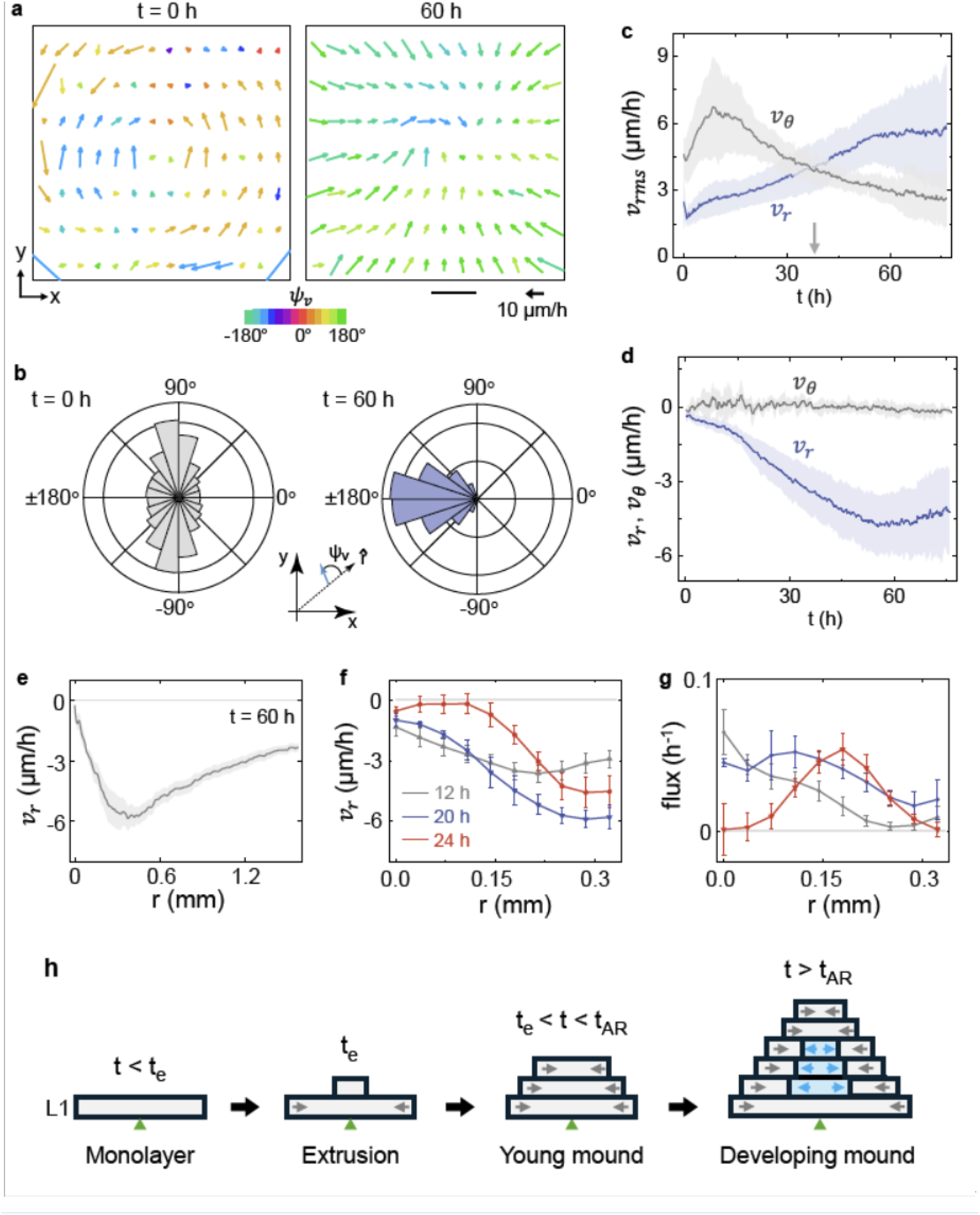
Dynamics of myoblast mound formation driven by a vortex defect. **a**, Representative velocity fields at confluency t = 0 h (left) and at time t = 60 h (right). Velocities were obtained by PIV from phase contrast movies (see Supplementary video 3 and 4). At 60 h, mounds are multilayered, and the PIV calculation integrates all layers. The color encodes the orientation of the velocities *ψ*_*v*_, with respect to the radial direction. Scale bar: 100 μm. **b**, Probability density functions (pdfs) of *ψ*_*v*_ at t = 0 h (grey) and t = 60 h (blue). The initial bidirectional azimuthal flow in the confluent monolayer gives way to a radial converging flow as the multilayers form and the mound matures. **c**, Time evolutions of the spatially averaged root-mean-square (rms) components of the velocity: *v*_*θ*_ (parallel to the direction of rails), and *v*_*r*_ (perpendicular to the direction of rails). The intensities of the two flows balance at t = t_AR_ ~ 38 ± 8 h (arrow). **d**, Time evolution of spatially averaged azimuthal *v*_*θ*_ and radial *v*_*r*_ velocity components. *v*_*θ*_ averages to 0 because the two rotation directions are equivalent. In contrast, *v*_*r*_ is negative and increases in absolute value as the flows converge unidirectionally faster and faster toward the center. **e**, Radial profile of the velocity component *v*_*r*_ at t = 60 h (negative values for converging flows). The position *r* = 0 corresponds to the vortex center. As time goes by, the center part of the mound becomes the locus of collective cell extrusion which shifts the position of highest flow away from the origin (here at ~ 400 μm from the origin). Velocities are measured from phase contrast images. **f**, Radial profiles of *v*_*r*_ in Layers 1 at t = 12 h (grey), t = 20 h (blue), t = 24 h (red). Note the different horizontal scale compared to panel **e**. Velocities measured by PIV from confocal images at z = 2 μm (Layer 1). **g**, Radial profiles of the cell flux from Layer 1 to Layer 2 at t = 12 h (grey), t = 20 h (blue) and t = 24 h(red). At t < *t*_*AR*_, cells mainly extrude near the defect site (r = 0 mm). For t < *t*_*AR*_, the position of the highest permeation flow shifts away from the defect site (see also Extended data Fig.7c). Velocities measured from confocal images at z = 2 μm (Layer 1). **h**, Schematic timeline of the process of cellular mound formation in the *x* − *z* plane. L1 is the first cell layer in direct contact with the substrate. t_e_ and t_AR_ are respectively the time at which the cells begin to form a bilayer and the time at which radial cell flow becomes dominant over azimuthal flow. Arrows: direction of the in-plane radial cell flows. Green triangle: Defect core position. With time, the convergent flow is maintained only around the mound while the center part becomes divergent. Permeation flows are not represented. Gray: vortex orientation. Blue: aster orientation. In panels c-d, data are presented as mean values +/-SD. In panels e-g, data are presented as mean values +/-SEM. Statistical information is provided in Extended data Table 1.

The convergent radial flow in Layer 1 became dominant over the fluctuating azimuthal flows at a crossover time t_AR_ = 38 ± 8 hours post-confluence (Figure 2c, Extended data Fig. 2b). Importantly, convergent flow extended over a spatial range of several millimeters (Figure 2e), with a maximum initially close to the core and shifting away from it as a second layer formed and spread over the initial monolayer (Figure 2f).

## Multilayering and formation of 3D mounds result from interlayer permeation

To investigate the dynamics of formation of the mounds, we used confocal live imaging between t = 12 h and t = 40 h. Even though the actual layers were intertwined (Fig. 2h), we describe layers as bidimensional nematic phase. This is consistent with the small mean elevation angle *ϕ* previously mentioned (see Extended data Fig. 3). In other words, we describe each layer as a bidimensional vortex phase. The first layer directly in contact with the substrate is referred to as Layer 1, and the subsequent layers are named Layer 2, …, Layer n. In each layer, we analyzed the radial component of the cell velocity, *v*_*r*_ in a 670 µm x 670 µm region centered at the defect core (Methods, Extended data Fig. 1j).

At t = t_e_ ~ 12 hours, myoblasts started to extrude at the defect core to form Layer 2, which expanded laterally on Layer 1, thereby forming a bilayer in which both layers took the vortex orientation of the substrate (Extended data Fig. 4). From t = 12 h to 32 h, the number of layers increased from 2 to 6, all of them being oriented as vortex (Extended data Fig. 7). All layers featured radial cell flows converging towards the core (*v*_*r*_ < 0) (Figure 2f, Extended data Fig. 7a). Of notice, when t > t_AR_, radial flows became slightly divergent (*v*_*r*_ > 0) close to the center of the middle layers (Figure 2h, Extended data Fig. 7b, Supplementary video 6). In this central region of divergent flows, cells gradually switched their orientation from vortex to aster (Figure 1g-i, and Extended data Fig. 7d-f, Supplementary video 6), while at the edge of the mound, a thick (~ 100 µm) “shell” remained where vortex orientation and its associated convergent flow persisted at a larger amplitude than the radial flows (Figure 1g-i, Extended data Fig.7a,b,d-f). In this shell, cells migrated along their short axis (Supplementary video 5), as already observed in the initial stages and in other bilayering processes [18]. By contrast, in the central part, cells migrated along their *long* axis (Extended data Fig. 7g, Supplementary video 6).

Assuming that the cell layers are incompressible and ignoring cell death or proliferation, a convergent 2D radial flow necessarily results in out-of-plane interlayer cell fluxes [37–39]. Indeed, exchange of cells between adjacent layers can be directly observed in experiments (Supplementary video 7). To account for this “*permeation*” normal to the layers, we include in the conservation equations the rate of cells extruding from a given layer and intercalating into an adjacent layer or into the external medium. For an incompressible stratified fluid, the velocity divergence of the i-th layer 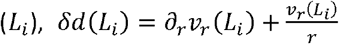 is related to the flux from *L*_*i*_ to, *L*_*i* + 1_, *j*(*i* → *i* + 1), and from *L*_*i* − 1_ to *L*_*i*_, *j*(*i* − *i* → 1), by the relation *δd*(*L*_*i*_) = − *j*(*i* → *i* + 1) + *j*(*i* − *i* → 1). As cells of Layer 1 extruded at the core and formed Layer 2 the process reiterated from Layer 2 to Layer 3, then Layer 4 etc. (Figure 2g-h, Extended data Fig.7). Permeation was initially maximal at the center, but the peak moved to larger *r* values at later times until reaching the aster-to-vortex boundary where the maximum of permeation eventually localized (Figure 2g, Extended data Fig. 7c, g).

## A theory of multilayering at a +1 vortex defect

Cell orientations and dynamical flows highlight the complexity of the processes at play in mound formation. In the following, we theoretically model two key steps of multilayering: we first study the onset of formation of these mounds by describing the formation of bilayers and then multilayers. Then, taking another approach, we describe the time-evolution of the 3D profiles of mature mounds. Only the main lines of our reasoning are developed here. The full calculations can be found in Supplementary Notes 1-2.

We first describe the bilayering observed at the core of the defect with a continuum description of stratified layers modeled as active nematic fluids with parameters coarse-grained at the tissue level. We first model the dynamics of a single layer, (Figure 3a and Supplementary Note 2A), and then, of a two-layer system (Figure 3e and Supplementary Note 2B). Layers are described as identical active nematic fluids where activity encompasses active force densities, such as gradients of active anisotropic stresses or nematic traction forces. Each layer is characterized by its in-plane velocity field and director field. Out-of-plane flows are modeled as interlayer permeation flows that are assumed to be proportional to the difference between the stress trace of successive stacked layers, or a layer and its external environment [37–41].

**Figure 3:**
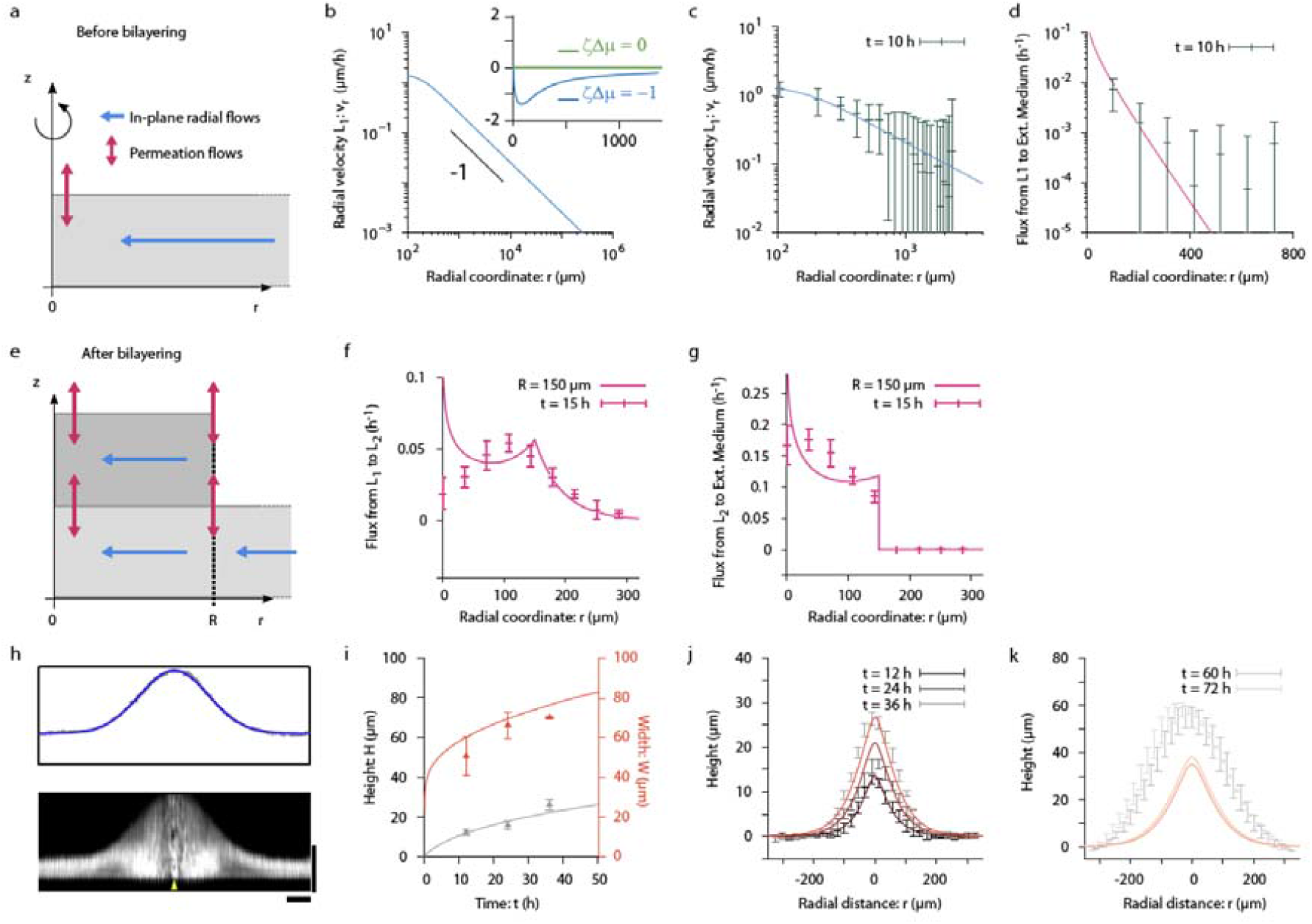
Modelling mound formation. **a**, Schematic of an active nematic monolayer plated on a vortex structure (the core of the defect is at the origin). Because the system is symmetric in rotation around the axis, only the plane defined by the axial coordinate and the radial coordinate are shown. In-plane radial flows are figured in Blue and out-of-plane permeation flows between a layer and the external environment in Magenta. **b**, Theoretical radial profile of the radial component of the velocity before the onset of bilayering (Eq. S48 in Supplementary Note 2A). Main panel: absolute value, Log-Log plot; inset: signed values for two values of the active anisotropic stress amplitude; all other active processes are set to zero. Lin-lin plot. Parameter values are given in Table S1 of the Supplementary Notes 2B. **c**, Comparison of the theoretical radial profile of the radial component of the velocity (absolute values) with experimental data in Layer1 (L1) in contact with the substrate, before the onset of bilayering. t = 10 h (green points). Solid curve: theoretical fit of Eq. S48 in Supplementary Note 2A on the experimental points. Parameter values in panels c and d are identical and are given in Table S1 of the Supplementary Notes 2B. **d**, Permeation flux from Layer 1 to external medium before the onset of bilayering. Points are experimental data (t = 10 h). Solid curve: theoretical fit of Eq. S49 in Supplementary Note 2A on these points. **e**, Schematic of an active nematic bilayer. Layer 1 is infinite, but Layer 2 has a finite area (radius *R*). See panel (a) for conventions. Magenta double-arrows indicate out-of-plane permeation flux across Layer 1 and Layer 2, and between Layer 2 and the external environment. R is the radius of Layer 2. **f**, Theoretical permeation flux from Layer 1 to Layer 2 in a bilayer system (solid line - Layer 2 is assumed to have an extension *R* = 150 μ*m*), and comparison with experimental data (points) at t = 15 h. Parameter set was chosen to qualitatively reproduce the experimental data. Parameter values in panels f and g are identical and are given in Table S1 of the Supplementary Notes 2B. **g**, Theoretical permeation flux from Layer 2 to external medium in a bilayer system (solid line - Layer 2 is assumed to have an extension *R* = 150 μ*m*), and comparison with experimental data at t = 15 h (points). Parameter set was chosen to qualitatively reproduce the experimental data. Parameter values are given in Table S1 of Supplementary Notes 2B. **h**, Angle-averaged profile of a representative mature cell mound (t = 65 h). Note the bell shape. The blue line is a gaussian fit of the experimental profile. Horizontal scale bar = 100 μm, vertical scale bar = 50 μm. Fixed mound, reconstruction from confocal microscopy, C2C12 β-actin–mCherry cells. **i**, Time evolution of the height (gray) and width (red) of cellular mounds in the first 40 h of their development (extracted from fits such as the one represented in panel (**h**), see Methods). Average over 4 experiments. Solid curves are theoretical fits by equations S74 and S76 in Supplementary Note 3A. **j**,**k**: Angle-averaged *x* − *z* profiles of a representative mound at different times as indicated in the legend. Solid lines are theoretical profiles for the parameter set *v*_*o*_ = 1 μ*m*/*h, ℓ* = 60 μ*m* and *χ*_0_/*v*_0_ *ℓ*_0_ 1, which is compatible with previous fits (panel (**i**)). Of notice, the good agreement at early times (**j**) and the deviation from the theoretical model as the mound ages (**k**). Velocities are measured from phase-contrast images in panels c and d, and from confocal images in panels f and g. In panels c-d, data are presented as mean values +/-SD. In panels f-g and i-k, data are presented as mean values +/-SEM.

Furthermore, based on our experimental observations, we make several assumptions: First, the director field in all layers is assumed to be parallel to the substrate plane. As confirmed by experimental observations (Extended data Fig. 3), there is no out-of-layer tilt. Then, layers are incompressible. Cell death and division typically occur on timescales of the order of 1 day [41], which is larger than the typical timescale found from out-of-plane cell fluxes, a few hours in our experiments. Therefore, fluid sources or sinks are ignored in the following. Finally, Layer 1 in contact with the substrate has infinite extension. We impose that the alignment of Layer 1 mirrors the vortex structure of the substrate. By contrast, Layer 2 has a finite radius *R*(*t*), and its alignment and velocity are dynamic.

The vortex geometry is characterized by a curvature decaying with the distance *r* to the center as 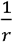. In an active nematic organized as a vortex, the interplay between active processes and this gradient of orientation results in an active long-range radial force density that varies also as 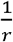 [22,42,43]. Its magnitude and direction are set by the values of the parameters of the active processes (Figure 3b inset, and Supplementary Note 2A). In the present case, parameters estimated from previous work yield a convergent force density [22] and yield steady-state converging flows also decaying as 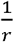, and thus long-range (Figure 3b, Supplementary Note 2A). This scaling with *r* is observed experimentally (Figure 3b-c, Extended data Fig. 8b). At the defect core compressive stresses are relieved by out of plane interlayer permeation (Figure 3d, Extended data Fig. 8, Extended data Fig. 8b).

We can define a characteristic length-scale, *ℓ*, resulting from a balance between interlayer dissipation due to permeation, dissipation due to substrate friction and intralayer viscous dissipation (Supplementary Notes 2A) [44]. Because of the *r* dependence of the radial flow, the permeation flow depends on the distance to the vortex core: At small distances ( *r* ≪ *ℓ*), it diverges logarithmically, while, at large distances (*r* ≫ *ℓ*), it vanishes exponentially. Therefore, unlike the in-plane radial flows, the permeation flow is short-range and localized near the vortex center.

We conclude that vortex oriented-myoblasts monolayers develop long-range convergent flows resulting eventually in the formation of multilayers at the core. From the position of the minimum of the radial flow in young mounds, we estimate *ℓ* ~ 150 μm (Figure 2f, t = 12 h).

After this onset of bilayering by a permeation mechanism, we can analyze the steady state flow profiles of the 2-layer system as a function of the radius *R* of Layer 2 (Figure 3e, Extended data Fig. 8c, and Supplementary Note 2B). For sufficiently small values of *R*, Layer 2 keeps a vortex orientation as experimentally observed. As detailed above for a single layer, the flux from Layer 1 to Layer 2 peaks at the vortex center. However, our analysis also shows another maximum at the edge of Layer 2, even though both layers share identical material parameters and the same director field configuration (Extended data Fig. 8d,e). These interlayer flows are measured experimentally (Figure 3f,g). They originate from differences in the stress trace between the two layers near the edge of Layer 2 (Supplementary Note 2B). As the flux from Layer 2 to the external medium (Figure 3g) peaks at the vortex center just as Layer 1 did initially (Figure 3d), a third layer subsequently forms by a similar mechanism.

## Diffusive-like dynamics of mature mounds

Although it is possible to describe the mound’s shape and its dynamics at long times (t > 30 h) by extrapolating the above-described permeation-based mechanism to many layers, this approach presents practical difficulties. Therefore, we develop here a simpler continuum model of the growth of such mature mounds (Supplementary Note 3). Our experiments show that the mounds are bell-shaped with a height and width that increase in time (Figure 1a,3h,i). Based on this evolution, we posit that the mound height evolves according to an effective diffusive dynamic with an effective diffusion constant *χ*_0_, and grows as a result of an influx of cells from Layer 1 to the mound at the defect core (Supplementary Note 3A). Of notice, we use the formalism of diffusion equations in the following for practical reasons, even though the physical mechanism of mound formation is not diffusive in the strict sense. This influx is described by the permeation flows from Layer 1 to the external medium (Supplementary Note 2A). This description captures the time evolution of the height and width of cell mounds from t = 12 h to t = 36 h (Figure 3i) with a single dimensionless parameter which corresponds to a rescaled diffusion coefficient: 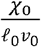, where *ℓ*_0_ is a screening length and *v*_0_ the magnitude of the velocity.

Fitting equations Eq. S74 and S76 in Supplementary Note 3A to the experimental values yields 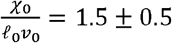 (Supplementary Note 3B). This value also describes well the entire mound shapes in this time span (Figure 3j). However, at later times (t > 60 h), the shape of the mounds deviates from the theory with mounds taller and wider compared to theoretical prediction (Figure 3k), suggesting that additional factors such as cell division, extra cellular matrix secretion, or time dependence of the scaled diffusion coefficient should be included in the calculation.

Altogether these results show that an effective diffusive dynamic encompassing out-of-plane permeation and radial variations of the in-plane flow reproduces the shape evolution of developing mounds.

## Activity drives mound formation independently of proliferation

Our analysis of 3D mound formation does not involve cell proliferation which contrasts with previous comparable experiments conducted with fibroblasts [20,21] where the increase of cell density at the core is clearly controlled by proliferation.

To confirm that proliferation is indeed not the driving force of mound formation in our experimental conditions (at least at early times), we performed experiments in which cell division was inhibited by Mitomycin C (MMC, see Methods). In these circumstances, we found indeed that myoblast mounds formed in the presence of MMC although they were slightly delayed in their development compared to the control (Figure 4). Notably, the complex cell organization associated with a nematic alignment in MMC-treated mature mounds was identical to the one observed in control experiments (Figure 4, Extended data Fig. 9). Let us note, however, that proliferation, even if not directly involved in mound formation, plays a very important role in setting the surface cell density. The delay in the formation of the mounds in the presence of MMC is likely to originate from a lower and constant cell density.

**Figure 4:**
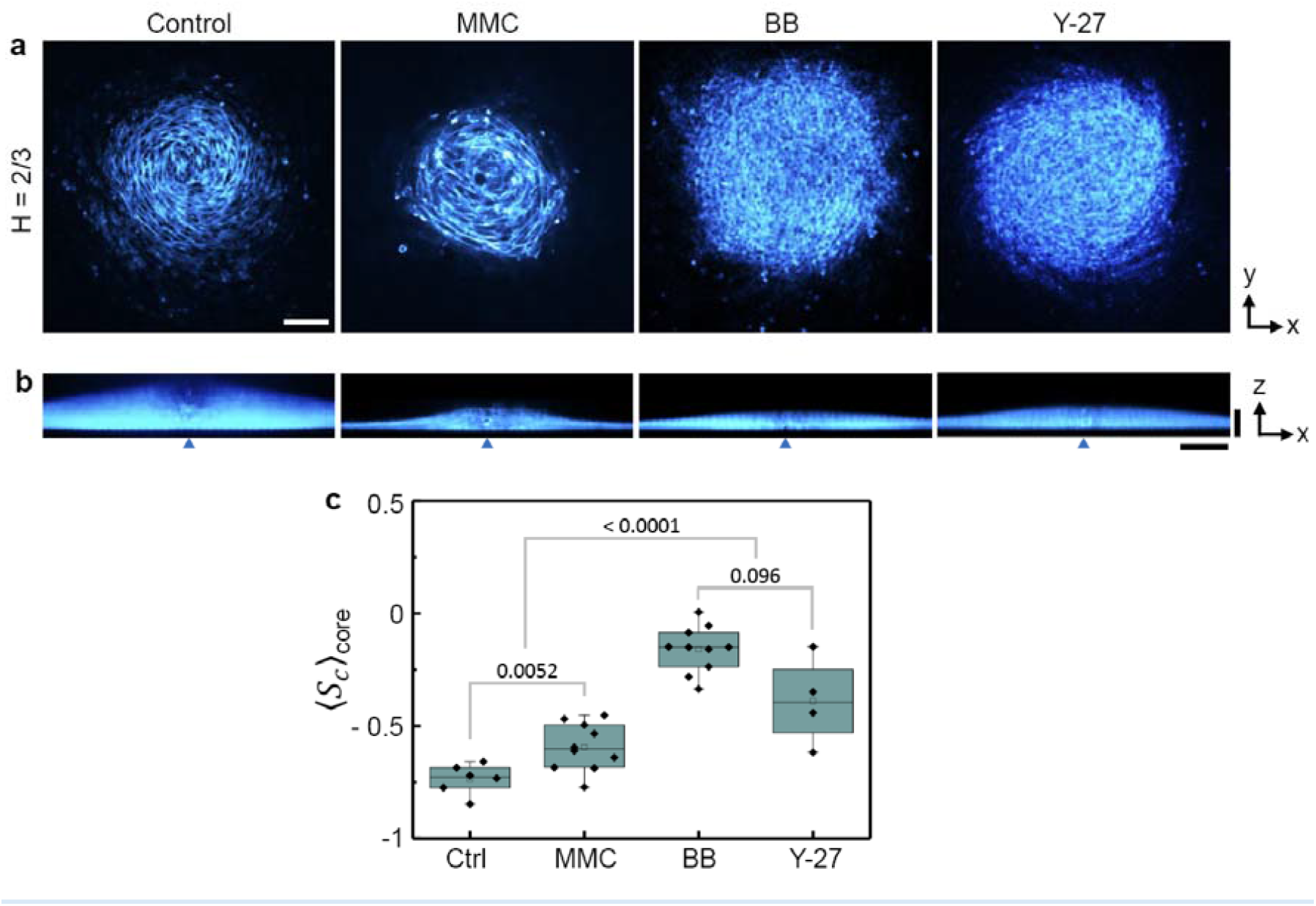
Effect of drug perturbations on cellular mound’s organization and growth. **a**, Representative confocal images of mature cell mounds at normalized height H = 2/3 after treatment with Mitomycin C (MMC) to inhibit proliferation, and with para-nitroblebbistatin (BB) and Y-27632 (Y-27) to inhibit cell contractility. Ctrl is DMSO. BB and Y27 effectively destroy cell alignment structures in the mound while MMC has a much lower impact. Therefore, myoblasts mounds are controlled by cell contractility and not proliferation (See also Extended data Fig. 9). **b**, Angle-averaged confocal images of mature cellular mounds along a diameter. The black triangles indicate the position of the defect core. **a, b**: Fixed cell mounds, C2C12 β-actin–mCherry cells, t = 110 h. Horizontal scale bar= 100 µm. **b**: vertical scale bar = 50 µm. **c**, Order parameter for in-plane cell orientation values after treatment, t = 110 h. Box plots with indication of the p-values. The mound structure is severely altered by inhibitors of contractility. Diamonds are experimental points (each point is an experiment). Error bars correspond to SDs over repeated runs. Box height represents the interquartile range (25th–75th percentile), with the horizontal bar indicating the median and the empty square representing the mean. p values are computed from a two-sample two-sided t-test. Statistical information is provided in Extended data Table 1.

In contrast, inhibiting cell contractility either with Blebbistatin or Y27-632 yielded flatter and disorganized cell mounds (Figure 4, Extended data Fig. 9). Our theory doesn’t directly predict the outcome of a decreased contractility on the structure of the mounds. Yet, in the absence of activity, our proposed mechanism of mound growth, including the vortex structure propagating from layer to layer, wouldn’t take place. We therefore conclude that activity and self-organization, but not proliferation, drive myoblast collective cell flows that shape the 3D mound at the defect center.

## Discussion

Altogether, we have shown that in-plane integer defects can orchestrate 2D morphogenetic flows in monolayers and eventually generate complex 3D tissue architectures encompassing precisely tuned cell orientations and a disclination line originating at the substrate vortex core and ending at the apex of the mound. The 3D dynamics of formation of these mounds is well captured by active matter theories including interlayer permeation but not cell proliferation.

The instrumental role of activity in the present system contrasts with previous studies in which cell accumulation at the defect center relied on cell proliferation [20,21]. This difference can be explained by the lower activity of the NIH-3T6 fibroblasts used in ref [21] compared to C2C12 myoblasts used here [32]. Note that the formation of a mound necessitates a source of cells. On our very wide substrates, this source can be the first monolayer itself over a long distance, with a small decrease in cell density corresponding to a thinning of the layer. Yet, cells continue to divide, and this proliferation term must be taken into consideration at late times.

Indeed, we can add cell division to the existing model by considering that within a cellular mound, new cells are produced at a constant rate, this rate being an additional fitting parameter. Even at this preliminary stage, this augmented model accounts very well for the entire trajectories, including late times. Moreover, the values of the parameters of the proliferation-free model (diffusion coefficient, length and time scales) are only weakly modified (i.e. similar order of magnitude but a factor 2 up or down) suggesting that cell division may indeed account for the late time regimes of experiments (Supplementary Notes 3C). We therefore conclude that two mechanisms can non-exclusively contribute to mound formation at topological defects. Weakly active cells rely on cell proliferation while more contractile cells use permeation hydrodynamic flows.

The analysis of the impact of the centripetal flow in Layer 1 on cells of Layer 2 allows to understand qualitatively the vortex-to-aster transition observed in the main part of developing mounds and the associated aster-vortex core-shell structure (Figure 1g-i). Interestingly, the orientation of the cells also impacts the distribution of the flow by a feedback mechanism. A detailed computation of this vortex-to-aster transition in the limiting case of a stack of two infinite layers is given in Supplementary Notes 2B.

Interestingly, this aster configuration of Layer 2 on top of the vortex-oriented Layer 1 leads to a perpendicular orientation of the two stacked layers. Such crisscross stacked muscle layers are common in organisms or organs that lack rigid skeletal elements [18,45,46] (hydrostats). Of notice, different mechanisms of formation of crisscross bilayers have been previously reported: i/ In monolayers of similar myoblasts in vitro, Layer 2 is initially oriented perpendicularly to Layer 1 as it spreads on it at a naturally occurring +1/2 defect [18]. ii/ Around the chick midgut, Layer 2 is initially disorganized and acquires its perpendicular orientation via the mechanical contractions of Layer 1 [45]. Finally, iii/ in the present work, Layer 2, initially oriented as Layer 1, progressively reorients perpendicular to it. Therefore, three distinct mechanisms generating crisscross bilayers have been identified so far, pointing to the importance of this structure, and making us speculate on a potential redundance and complementarity between them.

Finally, we propose that rationally designed surface micro-structures can be multiplexed to position and orient arrays of self-organized 3D cell ensembles. Myoblasts used in the present study were selected against differentiation, but applying this strategy to differentiating cells may provide ways based on self-organization to design arrays of tissue precursors or organoids.

## Acknowledgements

We dedicate this article to the memory of Caroline Giuglaris.

We thank the members of the Biology-inspired Physics at MesoScales (BiPMS) and in particular Mathilde Lacroix, and the Physical Approach to Biological Problems (PABP) group. The BiPMS group is a member of the Institut Pierre-Gilles de Gennes and has benefited from the technical contributions of the Unité d’Appui et de Recherche 3750 of the French National Centre for Scientific Research. We particularly thank Kevin Phan. The Molecular Biology and Cellular Biology platform of UMR168 is gratefully acknowledged and in particular A. Battistella and F. Cayrac for their help with the CRISPR transformation of the cells. We thank the Cell and Tissue Imaging core facility (PICT IBiSA), Institut Curie, member of the French National Research Infrastructure France-BioImaging (Grant No. ANR10-INBS-04). We particularly thank Olivier Renaud, Chloé Guedj, Lucie Sengmanivong, and Anne-Sophie Macé.

## Author contribution statement

HC and PS designed the research. HC and CG performed the experiments. HC, CBM, CG, and P.S. analysed the experimental data. CBM and JP developed the theoretical models. PS supervised the research. All authors contributed to writing the manuscript.

## Methods

### Cell culture and drug treatment

Experiments were conducted with wild type C2C12 myoblasts (provided by Clotilde Thery [Exosomes and Tumor Growth, INSERMU932, Institut Curie]) and C2C12 β-actin–mCherry cells (modified by the Curie BMBC facility using Crispr technique thanks to Aude Battistella and Fanny Cayrac) [47]. The C2C12 β-actin–mCherry cells were used for most of the experiments. Here, we used C2C12 myoblasts after a large number of passages after confluence (up to 40) to select them against differentiation [48,49]. Cells were cultured in standard DMEM medium (ThermoFisher Scientific) supplemented with 10% FBS (Dominique Dutscher) and 1% Penicillin–Streptomycin (Pen-Strep) mixture (Gibco) at 37°C under 5% CO_2_ partial pressure and 95% relative humidity. Under these conditions, division time was of the order of 1 day. For spinning disk confocal time-lapse experiments, we use phenol red-free HEPES DMEM (Gibco), high glucose, supplemented with 10% FBS, 1% Pen-Strep mixture. For inhibiting proliferation, cells were treated with Mitomycin-C (Merck) at 10 µM for 1 hour at 37°C. Then, cells were rinsed with balanced salt solution (PBS) which was replaced by fresh medium. For inhibiting contractility (myosin-II ATPase), cells were treated with para nitro-blebbistatin (Cayman Chemical) at 17 µM or with Y-27632 (Bertin Bioreagent) at 25 μM. Para nitro-blebbistatin is less phototoxic than blebbistatin and was therefore chosen for our experiments. For drug experiments, control samples were treated with the corresponding DMSO dilution.

### Microfabrication and cell seeding

Vortex defect (+1) microstructures (height of the rails=1.2µm, width 8 µm, period 16 µm) were fabricated by classical photolithography procedure through direct laser writing (µPG 101; Heidelberg instruments) or classical photolithography with UV illumination (MJB3 Mask aligner; Karl Süss). We spin-coated SU-8 photoresist (MicroChem) on a 4-inch silicon wafer, exposed, developed, and silanized these substrates with (Tridecafluoro-1,1,2,2-tetrahydrooctyl) trichlorosilane (abcr) for 2 hours. We then replicated these topographies in polydimethylsiloxane (PDMS, Sylgard 184 Dow Corning) by pouring a degassed mixture 10/1 w/w of the polymer and curing agent on the wafers, and curing overnight at 60°C.

Patterned PDMS slabs (also called indifferently in the text “vortex patterns” or “vortex microstructures”) (13 x 13 mm^2^) were then peeled, plasma-treated for 30 s, and placed in a 12-well glass bottom plate (CellVis). The PDMS slabs were then coated with a 10 µg/mL fibronectin solution (Fibronectin Bovine Protein, Bovine Plasma, Gibco) for 1 h. Cells were then plated on the microstructures at a density of approximately 175 000 cells/cm^2^, achieving confluence within 4 hours after seeding. Prior to imaging, cell monolayers were rinsed with PBS, and 5 mL of fresh medium was added. To identify cell orientation, some experiments were done by using mixtures of wild-type and 1-5% C2C12 β-actin–mCherry cells on defect microstructures coated with rhodamine-labeled fibronectin (EUROMEDEX) serving as a reference plane.

### Time-lapse microscopy

For time-lapse multi-field 2D experiments, PDMS slabs pre-patterned with a vortex defect are put in 12-well plates. Cells are then uniformly seeded all over the well (much wider than the structured PDMS). Phase-contrast and wide-field fluorescence images were acquired using an automated Olympus X71 inverted microscope (x10 objective) equipped with temperature, humidity and CO_2_ controls (Life Imaging Services). Image acquisitions were performed with a sCMOS camera (PrimeBSI, Teledyne) controlled by Metamorph (Universal Imaging). Images were acquired typically every 10 minutes. The field of view (FOV) was 1.3 x 1.3 mm^2^.

Time–lapse 3D imaging was acquired on spinning-disk confocal microscopes with x20 oil/water objectives (inverted Nikon Eclipse Ti-E with spinning disk CSU-W1/FRAP module or Nikon Eclipse Ti–2 with Yokogawa W1 spinning disk unit). For time-lapse spinning disk confocal live imaging, the FOV is 0.67 x 0.67 mm^2^. z-step = 2 µm. 3D image stacks were acquired every 20–30 minutes.

### Cell fixing and staining

Cells were fixed with 4% (wt/vol) paraformaldehyde (Electron Microscopy Sciences), permeabilized with Triton X-100 (EUROMEDEX), and saturated with 0.2% Bovine Serum Albumin (Sigma) and 2.5 % Normal Goat Serum (Invitrogen) in PBS. After fixation, filamentous actin was labelled by phalloidin-Tritc (Sigma-Aldrich, Cat. No. P1951, dilution 1:150), and cell nuclei were labelled with DAPI (Sigma, dilution: 1:1000). For Collagen IV staining, fixed cells were incubated with primary antibodies (Rabbit polyclonal anti–Collagen IV (Abcam, ab6586, 1:100 dilution)) for 2 days at 4°C and secondary antibodies (Thermo Fisher, Alexa Fluor® Chicken anti-Rabbit IgG (H+L), 1:100 dilution) for 30 min at room temperature. Fixed and stained samples were imaged by inverted spinning disk confocal microscopes (Nikon). z-step = 1 µm.

### Image analysis and data analysis

Analysis of the images was performed using Fiji public domain software [50] or Matlab (MatWorks). MATLAB R2021a was used for obtaining velocity fields by particle image velocimetry (PIV) with the Matlab toolbox PIVlab 2.46 [51]. The window size was set to 16 pixels (20.8 µm) with a 50 % overlap. A sliding-time average over 30 minutes was then applied. For phase contrast and fixed experiments, stitching of the FOVs was performed using the Grid/Collection stitching plugin [52,53], and the dimensions of the stitched images were typically 3 x 3 mm^2^. Spinning disk confocal images were not stitched (FOV= 0.67 x 0.67 mm^2^).

For characterizing velocity fields, the angle *ψ*_*v*_ is the angle between the orientation of local velocity *v*, obtained through PIV and their corresponding radial direction. *ψ*_*v*_ = ±90° denotes bi-directional azimuthal flows and *ψ*_*v*_ = −180° (resp. *ψ*_*v*_ = +180°) corresponds to inward (resp. outward) radial flows. To measure cell orientation, we analyzed fluorescent images of actin-labeled cells with the Fiji plugin OrientationJ [54]. The angle *ψ*_*c*_ is the angle between the cell orientation and radial direction *r* pointing away from the defect core, where the subscript *c* stands for cells. The director field being invariant to the transformation ***n*** → −***n***, the values of *ψ*_*c*_ and *ψ*_*c*_ + *π* are equivalent. Therefore, we restrict the range of values of *ψ*_*c*_ to 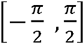. See Extended data Fig. 1 for schematics.

The order parameters were defined as *S*_*v*_ = *cos*(2*ψ*_*v*_) (resp. *S*_*c*_ = *cos*(2*ψ*_*v*_)), where the subscript *v* (resp. *c*) stands for velocity (resp. cells). *S* = −1 refers to vortex configuration, where velocities or cellular orientation directors are perpendicular to *r* (alignment in the direction of rails). *S* =+1 refers to aster configurations, where the velocity or cellular orientation is parallel to *r* (angle perpendicular to the direction of rails). In Figure 1e, ⟨*S*_*c*_ ⟩_*core*_ is an average of *S*_*c*_ within the region centered at the defect core, with size 0.15 mm^2^. Rescaling of z by mound height (Figure 1e and Extended data Fig. 6) was performed over different mounds (different experiments, different times (t > 40h)), after rescaling heights to 1.

The nematic order parameter, *Q* (Figure 1j), was computed as 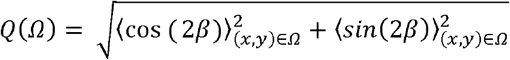, where the average is taken over a window *Ω* = 32.5 μm x 32.5 μm [32]. The angle *β* is defined as the angle of the director field *n* = (*cos* (*β*),*sin*(*β*) with respect to an arbitrary fixed x direction, See Extended data Fig. 1i. The procedure above was repeated for each z-plane to map *Q* in Figure 1j.

To obtain the cellular mound height and width in Figure 3i, each average height profile was fitted by a Gaussian function 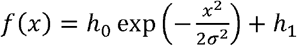 whose height *h*_0_ and width *σ* are fitting parameters and *h*_1_ = 6 μ*m* is the thickness of Layer 1. The procedure to fit the time evolution of the height and width assuming effective diffusive dynamics is detailed in Supplementary Notes 3B.

The shape index of the cells is defined as 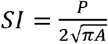 where P is the perimeter and A is the area of the cell. SI = +1 for a circular isotropic cell and SI > 1 for a spindle-shape cell.

Data analysis was performed with MATLAB (Mathworks) or with Origin (Originlab).

### Statistics

Experiments were conducted in separate wells of a glass-bottom 12-well plate, with each well representing an independent experiment. At least two replica were performed for each condition and results were pooled and averaged from all experiments acquired in these independent replicas. The number of analyzed wells, n, and the number of replica, N, are given in Extended data Table 1. Our sample sizes are similar or larger than those reported in previous publications [27,36].

## Data availability

The data that support the findings of this study and that are not in the paper, its extended data files, or in its supplementary information file, have been deposited in zenodo.org (DOI: 10.5281/zenodo.20612726).

## References

1. Kindberg, A., Hu, J. K. & Bush, J. O. Forced to communicate: Integration of mechanical andbiochemical signaling in morphogenesis. Curr. Opin. Cell Biol. 66, 59–68 (2020).

2. Chan, C. J.Heisenberg, C.-P. & Hiiragi, T. Coordination of Morphogenesis and Cell-Fate Specification in Development. Curr. Biol. 27, R1024–R1035 (2017).

3. Urdy, S. On the evolution of morphogenetic models: mechano-chemical interactions and an integrated view of cell differentiation, growth, pattern formation and morphogenesis. Biol. Rev. 87, 786–803 (2012).

4. Lecuit, T. & Le Goff, L. Orchestrating size and shape during morphogenesis. Nature 450, 189–192 (2007).

5. Friedl, P. & Gilmour, D. Collective cell migration in morphogenesis, regeneration and cancer. Nat. Rev. Mol. Cell Bio.l 10, 445–457 (2009).

6. Bertet, C., Sulak, L. & Lecuit, T. Myosin-dependent junction remodelling controls planar cell intercalation and axis elongation. Nature 429, 667–671 (2004).

7. Heisenberg, C.-P. & Bellaïche, Y. Forces in tissue morphogenesis and patterning. Cell 153, 948–962 (2013).

8. Le Verge-Serandour, M. & Turlier, H., Blastocoel morphogenesis: A biophysics perspective. Semin. Cell Dev. Biol. 130, 12–23 (2022).

9. Osterfield, M., Du, X., Schüpbach, T., Wieschaus, E. & Shvartsman, S. Y. Three-Dimensional Epithelial Morphogenesis in the Developing Drosophila Egg. Dev. Cell 24, 400–410 (2013).

10. Davidson, L. A. Epithelial machines that shape the embryo. Trends in Cell Biol. 22, 82–87 (2012).

11. Yamada, K. M. & Cukierman, E. Modeling tissue morphogenesis and cancer in 3D. Cell 130, 601–610 (2007).

12. Doostmohammadi, A., Ignés-Mullol, J., Yeomans, J. M. & Sagués, F. Active nematics. Nat. Commun. 9, 3246 (2018).

13. Chen, T., Saw, T. B., Mège, R.-M. & Ladoux, B. Mechanical forces in cell monolayers. J. Cell Sci. 131, jcs218156 (2018).

14. Doostmohammadi, A. & Ladoux, B. Physics of liquid crystals in cell biology. Trends in Cell Biol. 32, 140–150 (2022).

15. Duclos, G., Garcia, S., Yevick, H. G. & Silberzan, P. Perfect nematic order in confined monolayers of spindle-shaped cells. Soft Matter 10, 2346–2353 (2014).

16. Saw, T. B. et al. Topological defects in epithelia govern cell death and extrusion. Nature 544, 212–216 (2017).

17. Kawaguchi, K., Kageyama, R. & Sano, M. Topological defects control collective dynamics in neural progenitor cell cultures. Nature 545, 327–331 (2017).

18. Sarkar, T. et al. Crisscross multilayering of cell sheets. PNAS Nexus 2, pgad034 (2023).

19. Elsdale, T. & Foley, R. Morphogenetic aspects of multilayering in Petri dish cultures of human fetal lung fibroblasts. J. Cell Biol. 41, 298–311 (1969).

20. Endresen, K. D., Kim, M., Pittman, M., Chen, Y. & Serra, F. Topological defects of integer charge in cell monolayers. Soft Matter 17, 5878–5887 (2021).

21. Kaiyrbekov, K. et al. Migration and division in cell monolayers on substrates with topological defects. Proc. Natl. Acad. Sci. U.S.A. 120, e2301197120 (2023).

22. Zhao, Z. et al. Integer topological defects offer a methodology to quantify and classify active cell monolayers. Nat. Commun. 16, 2452 (2025).

23. Sanchez-Corrales, Y. E. & Blanchard, G. B. Radially patterned cell behaviours during tube budding from an epithelium. eLIFE 7, e35717, (2018).

24. Guruciaga, P. C., Ichikawa, T., Hiiragi, T. & Erzberger, A. Boundary geometry controls a topological defect transition that determines lumen nucleation in embryonic development. arXiv preprint arXiv:2403.08710 (2024).

25. Maroudas-Sacks, Y. et al. Topological defects in the nematic order of actin fibres as organization centres of Hydra morphogenesis. Nat. Phys. 17, 251–259 (2021).

26. Ravichandran, Y., Vogg, M., Kruse, K., Pearce, D. J. G. & Roux, A. Topology changes of Hydra define actin orientation defects as organizers of morphogenesis. Sci. Adv. 11, eadr9855 (2025).

27. Guillamat, P., Blanch-Mercader, C., Pernollet, G., Kruse, K. & Roux, A. Integer topological defects organize stresses driving tissue morphogenesis. Nat. Mater. 21, 588–597 (2022).

28. Li, M. et al. Regulation of an osteon-like concentric microgrooved surface on osteogenesis and osteoclastogenesis. Biomaterials 216, 119269 (2019).

29. Kepes, J. J. Cellular whorls in brain tumors other than meningiomas. Cancer 37, 2232–2237 (1976).

30. Zhang, J., Yang, N., Kreeger, P. K. & Notbohm, J. Topological defects in the mesothelium suppress ovarian cancer cell clearance. APL Bioeng. 5, 036103 (2021).

31. Szporn, A. H., Yuan, S., Wu, M. & Burstein, D. E. Cellular swirls in fine needle aspirates of papillary thyroid carcinoma: a new diagnostic criterion. Mod. Pathol. 19, 1470–1473 (2006).

32. Duclos, G., Erlenkämper, C.Joanny, J.-F. & Silberzan, P. Topological defects in confined populations of spindle-shaped cells. Nat. Phys. 13, 58–62 (2017).

33. Borshch, V. et al. Nematic twist-bend phase with nanoscale modulation of molecular orientation. Nat. Commun. 4, 2635 (2013).

34. Nguyen, A. T., Sathe, S. R. & Yim, E. K. F. From nano to micro: topographical scale and its impact on cell adhesion, morphology and contact guidance. J. Phys.: Condens. Matter 28, 183001 (2016).

35. Leclech, C., Gonzalez-Rodriguez, D., Villedieu, A. et al. Topography-induced large-scale antiparallel collective migration in vascular endothelium. Nat Commun 13, 2797 (2022).

36. Lacroix, M. et al. Emergence of bidirectional cell laning from collective contact guidance. Nat. Phys. 20, 1324–1331 (2024).

37. Heslot, F., Fraysse, N. & Cazabat, A. M. Molecular layering in the spreading of wetting liquid drops. Nature 338, 640–642 (1989).

38. Helfrich, W. Elastic properties of lipid bilayers: Theory and possible experiments. Phys. Rev. Lett. 23, 372–375 (1969).

39. de Gennes, P. G. & Cazabat, A. M. Etalement d’une goutte stratifiée incompressible. C. R. Acad. Sci. Paris, Ser. II 310, 1601–1606 (1990).

40. Beaune, G. et al. How cells flow in the spreading of cellular aggregates. Proc. Natl. Acad. Sci. U.S.A. 111, 8055–8060 (2014).

41. Humphrey, G. W. et al. GREG cells, a dysferlin-deficient myogenic mouse cell line. Exp. Cell Res. 318, 127–135 (2012).

42. Kruse, K., Joanny, J. F., Jülicher, F., Prost, J. & Sekimoto, K. Asters, Vortices, and Rotating Spirals in Active Gels of Polar Filaments. Phys. Rev. Lett. 92, 078101 (2004).

43. Blanch-Mercader, C., Guillamat, P., Roux, A. & Kruse, K. Integer topological defects of cell monolayers: mechanics and flows. Phys. Rev. E 103, 012405 (2021).

44. Ranft, J. et al. Tissue dynamics with permeation. Eur. Phys. J. E 35, 46 (2012).

45. Huycke, T. R. et al. Genetic and Mechanical Regulation of Intestinal Smooth Muscle Development. Cell 179, 90–105.e21 (2019).

46. Gilbert, R. J., Napadow, V. J., Gaige, T. A. & Wedeen, V. J. Anatomical basis of lingual hydrostatic deformation. J. Exp. Biol. 210, 4069–4082 (2007).

## Methods-only references

47. Lyu, M. et al. Efficient CRISPR/Cas9-mediated gene editing in mammalian cells by the novel selectable traffic light reporters. Int. J. Biol. Macromol. 243, 124926 (2023).

48. Sharples, A. P., Al-Shanti, N., Lewis, M. P. & Stewart, C. E. Reduction of myoblast differentiation following multiple population doublings in mouse C2C12 cells: a model to investigate ageing? J. Cell. Biochem. 112, 3773–3785 (2011).

49. Shahini, A. et al. NANOG restores the impaired myogenic differentiation potential of skeletal myoblasts after multiple population doublings. Stem Cell Res. 26, 55–66 (2018).

50. Schindelin, J. et al. Fiji: an open-source platform for biological-image analysis. Nat. Methods 9, 676–682 (2012).

51. Thielicke, W. & Stamhuis, E. J. PIVlab – Towards User-friendly, Affordable and Accurate Digital Particle Image Velocimetry in MATLAB. J. Open Res. Softw. 2, e30, (2014).

52. Schneider, C. A., Rasband, W. S. & Eliceiri, K. W. NIH Image to ImageJ: 25 years of image analysis. Nat. Methods 9, 671–675 (2012).

53. Preibisch, S., Saalfeld, S. & Tomancak, P. Globally optimal stitching of tiled 3D microscopic image acquisitions. Bioinformatics 25, 1463–1465 (2009).

54. Püspöki, Z., Storath, M., Sage, D. & Unser, M. Transforms and operators for directional bioimage analysis: a survey. In Focus on Bio-Image Informatics Vol. 219 (eds De Vos, W. H., Munck, S. & Timmermans, J.-P.) 69–93 (Springer, 2016).

